# P_*AOX1*_ expression in mixed-substrate continuous cultures of *Komagataella phaffii* (*Pichia pastoris*) is completely determined by methanol consumption regardless of the secondary carbon source

**DOI:** 10.1101/2021.09.12.459941

**Authors:** Anamika Singh, Atul Narang

## Abstract

The expression of recombinant proteins by the *AOX1* promoter of *Komagataella phaffii* is typically induced by adding methanol to the cultivation medium. Since growth on methanol imposes a high oxygen demand, the medium is often supplemented with an additional secondary carbon source which serves to reduce the consumption of methanol, and hence, oxygen. Early research recommended the use of glycerol as the secondary carbon source, but more recent studies recommend the use of sorbitol because glycerol represses P_*AOX1*_ expression. To assess the validity of this recommendation, we measured the steady state concentrations of biomass, residual methanol, and LacZ expressed from P_*AOX1*_ over a wide range of dilution rates (0.02–0.20 h^−1^) in continuous cultures of the Mut^+^ strain fed with methanol + glycerol (repressing) and methanol + sorbitol (non-repressing). We find that under these conditions, the specific P_*AOX1*_ expression rate is completely determined by the specific methanol consumption rate regardless of the type (repressing/non-repressing) of the secondary carbon source. In both cultures, the specific P_*AOX1*_ expression rate is proportional to the specific methanol consumption rate provided that the latter is below 0.15 g/(gdw-h); beyond this threshold consumption rate, the specific P_*AOX1*_ expression rate of both cultures saturates to the same value. Analysis of the data in the literature shows that the same phenomenon also occurs in continuous cultures of *Escherichia coli* fed with mixtures of lactose plus repressing/non-repressing carbon sources. The specific P_*lac*_ expression rate is completely determined by the specific lactose consumption rate regardless of the type of secondary carbon source, glycerol or glucose.

**Key points:**

- P_*AOX1*_ expression rate is completely determined by the methanol consumption rate.
- Sorbitol is not necessarily superior secondary carbon source than glycerol.

## Introduction

The methylotrophic yeast *Komagataella phaffii*, referred to earlier as *Pichia pastoris* (Kurtzman, 2005; Kurtzman, 2009), is a popular expression host (Schwarzhans *et al*., 2016). There are several reasons for this, but the most important one is that *K. phaffii* has an unusually strong and tightly regulated promoter which drives the expression of alcohol oxidase (AOX) in the presence of methanol (Higgins and Cregg, 1998; Ahmad *et al*., 2014; Gasser and Mattanovich, 2018). To be sure, *K. phaffii* has two alcohol oxidase genes, *AOX1* and *AOX2*, with corresponding promoters, P_*AOX1*_ and P_*AOX2*_, but P_*AOX1*_ is used to drive recombinant protein expression since it is ~10 times stronger than P_*AOX2*_ (Cregg *et al*., 1989).

In the first expression system constructed with *K. phaffii*, the wild-type strain was used as host, and recombinant protein was expressed under the control of P_*AOX1*_ by using methanol as inducer (Cregg *et al*., 1985). Although this Mut^+^ (methanol utilization plus) strain yielded excellent recombinant protein expression, the use of methanol as inducer led to several operational problems (McCauley-Patrick *et al*., 2005; Cos *et al*., 2006; Jahic *et al*., 2006; Jungo *et al*., 2007a; Arnau *et al*., 2011; Potvin *et al*., 2012; Yang and Zhang, 2018; García-Ortega *et al*., 2019; Liu *et al*., 2019). Indeed, methanol is inflammable which poses safety issues. Moreover, methanol metabolism results in high oxygen demand and heat generation, as well as excretion of toxic metabolites such as formaldehyde that inhibit growth (Jungo *et al*., 2007b; Juturu and Wu, 2018).

The problems stemming from the use of methanol as inducer led to several strategies for reducing methanol consumption. One strategy was to engineer the host strain by deleting either *AOX1* or both *AOX1* and *AOX2*, thus producing the Mut^s^ (methanol utilization slow) and Mut^s^ (methanol utilization minus) strains, respectively, whose capacity to consume methanol is substantially impaired or abolished (Chiruvolu *et al*., 1997). Another strategy was to introduce into the medium, in addition to the *primary* or *inducing* carbon source methanol, a *secondary* or *non-inducing* carbon source that supports growth but not induction. This reduces methanol consumption due to the sparing effect of the secondary carbon source, and increases the volumetric productivity due to the enhanced cell growth derived from metabolism of the secondary carbon source (Brierley *et al*., 1990; Egli and Mason, 1991, Jungo *et al*., 2007a; Jungo *et al*., 2007b; Paulova *et al*., 2012).

The foregoing strategies have led to reduced methanol consumption, but they can also result in decreased recombinant protein expression. Recently, we found that host strain engineering decreases recombinant protein expression substantially — the specific productivities of the engineered Mut^s^ and Mut^−^ strains are respectively 5- and 10-fold lower than that of the Mut^+^ strain (Singh and Narang, 2020). Since these three strains differ only with respect to their capacity for methanol consumption, the methanol consumption rate is an important determinant of the P_*AOX1*_ expression rate.

The goal of this work is to quantify the extent to which P_*AOX1*_ expression is affected by addition of a secondary carbon source to the medium. It is commonly held that this is determined by the type of the secondary carbon source. Specifically, these carbon sources have been classified as *repressing* or *non-repressing* based on the P_*AOX1*_ expression levels observed in *batch* cultures of the Mut^−^ strain grown on mixtures of methanol and various secondary carbon sources (Inan and Meagher, 2001). Repressing carbon sources, such as glycerol, abolish P_*AOX1*_ expression, whereas non-repressing carbon sources, such as sorbitol, permit P_*AOX1*_ expression. The same conclusion has been reached from studies of mixed-substrate growth in fed-batch cultures (Brierley *et al*., 1990; Thorpe *et al*., 1999; Xie *et al*., 2005; Çelik *et al*., 2009; Wang *et al*., 2010; Gao *et al*., 2012; Niu *et al*., 2013; Carly *et al*., 2016; Azadi *et al*., 2017; Chen *et al*., 2017) and continuous cultures (Jungo *et al*., 2006; Jungo *et al*., 2007a; Jungo *et al*., 2007b; Canales *et al*., 2015; Berrios *et al*., 2017). Indeed, even though glycerol is commonly used as the secondary carbon source, the use of sorbitol has been almost unanimously recommended on the grounds that glycerol represses P_*AOX1*_ expression.

Most of the comparative studies cited above used constant fed-batch cultures, but these data can be difficult to interpret physiologically because the specific growth rate decreases throughout the course of the experiment (Nieto-Taype *et al*., 2020). The comparative studies with continuous cultures are reviewed at length in the Discussion. Here, it suffices to note that many of these studies were performed at a fixed dilution rate *D*, and hence, specific growth rate (Jungo *et al*., 2007a; Jungo *et al*., 2007b; Berrios *et al*., 2017). We reasoned that comparative studies over a wide range of *D* could yield deeper physiological insights into the factors governing P_*AOX1*_ expression. Moreover, the optimal operating conditions determined in continuous cultures can also inform optimal protein production in exponential fed-batch cultures (Jungo *et al*., 2007a; Jungo *et al*., 2007b).

We were therefore led to study P_*AOX1*_ expression in continuous cultures of *K. phaffii* operated at various dilution rates with fixed concentrations of methanol + glycerol and methanol + sorbitol. To this end, we used a Mut^+^ strain expressing LacZ from P_*AOX1*_, but we also measured the AOX level to check the consistency of the data. We find that the specific P_*AOX1*_ expression rate is completely determined by the specific methanol consumption rate regardless of the type (repressing/non-repressing) of the secondary carbon source.

## Materials and Methods

### Microorganism and growth medium

A *K. phaffii* Mut^+^ strain, GS115 (*his4*) was procured from J. M. Cregg, Keck Graduate Institute, Claremont, CA, USA and was genetically modified to express a recombinant β-galactosidase protein. Details of the strain construction have been presented elsewhere (Singh and Narang, 2020). The resulting strain was called Mut^+^ (pSAOH5-T1) and was used for this study. Stock cultures were stored in 25% glycerol at –80 °C.

The minimal medium composition used for shake-flask as well as chemostat cultivations was chosen such as to ensure stoichiometric limitation by the carbon and energy sources as described in Egli and Fiechter (1981). The defined medium was supplemented with either glycerol (~3.1 g |^−1^) or a mixture of methanol (~1.6 g |^−1^)/(~3.2 g |^−1^). and glycerol/sorbitol (~1.5 g |^−1^) as carbon sources and in addition, contained 100 mM phosphate buffer (pH 5.5), 15.26 g NH_4_Cl, 1.18 g MgSO_4_·7H_2_O, 110 mg CaCl_2_·2H_2_O, 45.61 mg FeCl_3_, 28 mg MnSO_4_·H_2_O, 44 mg ZnSO_4_·7H_2_O, 8 mg CuSO_4_·5H_2_O, 8.57 mg CoCl_2_·6H_2_O, 6 mg Na_2_MoO_4_·2H_2_O, 8 mg H_3_BO_3_, 1.2 mg KI, 370 mg EDTA disodium salt, 2.4 mg biotin per liter. All components of the defined medium were prepared and sterilised by either filtration or autoclaving as separate stock solutions and then mixed before cultivation.

### Inoculum preparation and chemostat cultivation

When required, cells were revived in a 100 mL shake flask containing 10 mL minimal medium supplemented with a suitable carbon source at 30 °C and 200 rpm. These primary cultures were sub-cultured once before inoculating the reactor precultures (in the same cultivation medium as prepared for the reactor vessel) which were then used as an inoculum for the bioreactor.

Chemostat cultivations were performed using bench-scale 0.5 L mini bioreactors modified to support chemostat operation and equipped with pH, DO, temperature, level and agitation controls (Applikon Biotechnology, The Netherlands) at working volumes of 0.3 L. The cultivation temperature was always maintained at 30 °C and pH at 5.5 by the automatic addition of 2 M NaOH. An integrated mass flow controller ensured a constant supply of air to the reactor vessel at 80 mL min^−1^. Dissolved oxygen levels were monitored by a polarographic probe calibrated with respect to an air-saturated medium. Cultures were agitated to ensure fast mixing as well as aerobic conditions such that the DO level always remained above 60 %. A silicone based anti-foam agent was added to the reactor vessel as and when required to prevent foam formation and wall growth. For chemostat mode operation, the dilution rate was set by fixing the input feed flow rate while a constant volume was maintained inside the reactor vessel by controlling the output feed flow rate via proportional control based on the on-line monitoring of the change in weight of the reactor vessel. The O_2_ and CO_2_ levels in the off-gas were measured using a Tandem gas analyser (Magellan Biotech, UK). After inoculation, cells were grown in batch phase for some time to allow exhaustion of the initial carbon source (indicated by a rise in DO level), followed by initiating the input and output feed supplies. At any particular dilution rate, steady-state samples were withdrawn after 5-6 liquid residence times. In general, three samples were collected for each dilution rate, separated by an interval of one liquid residence time.

### Sample collection and processing

For determination of residual substrate concentration inside the reactor, samples were withdrawn directly from the vessel. To achieve rapid biomass separation, culture samples were withdrawn using vacuum through a sampling tube attached to a 0.2-micron syringe filter and stored at –20 °C until analysis. Samples for determination of biomass and enzyme activities were collected in a sampling bottle kept on ice. Biomass samples were processed immediately, while samples for measuring enzyme activities were pelleted, washed and stored at –20 °C until processing.

### Substrate analysis

Glycerol and sorbitol concentrations were estimated by high-performance liquid chromatography (HPLC) analysis (1100 series, Agilent Technologies, Palo Alto, USA) with detection limits of ~1 mg/l and ~30 mg/l. An ion-exclusion chromatography column from Phenomenex, California, USA (ROA-Organic acid H^+^ column, 300 × 7.8 mm, 8 μm particle size, 8% cross linkage) with a guard column (Carbo-H cartridges) was used with 5 mM H_2_SO_4_ in ultrapure water as mobile phase supplied at a constant flow rate of 0.5 mL min^−1^. The column chamber was maintained at 60 °C and a refractive index detector was used for substrate measurement. Methanol concentrations were determined with a gas chromatograph equipped with a flame ionisation detector (GC-FID) (7890A, Agilent Technologies, Palo Alto, USA) using a HP-PLOT/Q column (30 m × 0.32 mm, 20 μm) from Agilent Technologies and nitrogen as the carrier gas. The detection limit for methanol was ~5 mg/l.

### Dry cell weight measurement

A known volume of the fermentation broth was collected and pelleted in a pre-weighed centrifuge tube. Pellets were washed twice with distilled water and then dried at 80 °C to constant weight.

### Cell-free extract preparation

Culture samples were collected on ice and immediately centrifuged at 4 °C to collect cells. The cell pellets were washed twice with phosphate buffer (100 mM, pH 7.4) and stored at –20 °C until analysis. For cell lysis, pellets were resuspended in 100 μl of chilled breaking buffer (Jungo *et al*., 2006). Acid-washed glass beads (0.40–0.45 mm diameter) were added to the resulting slurry followed by alternate vortexing (1 min) and resting (on ice for 1 min) steps. This cycle was repeated 4-5 times, after which the cell debris was removed by centrifugation. Cell-free extracts (supernatant) were collected in fresh tubes kept on ice and immediately used for the estimation of enzyme activities. The Bradford assay was used for the estimation of the total protein content of the cell-free extracts for which bovine serum albumin served as standard (Bradford 1976).

### β-galactosidase assay

β-galactosidase assays were performed according to the method described by Miller (1972) with modifications. Briefly, cell-free extracts were appropriately diluted and mixed with Z-buffer containing β-mercaptoethanol (Miller 1972) and incubated at 30 °C in a water-bath for 15-20 minutes. The reaction was started by adding ONPG and stopped by adding Na_2_CO_3_ when sufficient colour had developed. The specific β-galactosidase activity was calculated with the formula

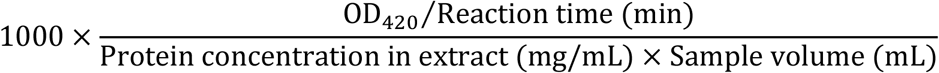

and expressed in units mgp^−1^ where mgp denotes mg of total protein.

### Alcohol oxidase assay

Appropriate dilutions of the cell-free extracts were used to measure alcohol oxidase activities based on the method adapted from Jungo et al (2006). A fresh 2x stock of the assay reaction mixture containing 0.8 mM 4-aminoantipyrine, 50 mM phenolsulfonic acid, freshly prepared 4 U/mL horseradish peroxidase in potassium phosphate buffer (200 mM, pH 7.4) was prepared before setting up the assays. 100 μL of the diluted cell-free extracts were mixed with 25 μL methanol and incubated at 30 °C for 10 minutes. After this, 100 μL of the 2x reaction mixture stock was added to the mix at time t = 0 to start the reaction and the increase in absorbance at 500 nm was monitored every 30 seconds for 10 minutes using a microplate reader (SpectraMax M2e, Molecular Devices Corporation, CA, USA). The specific alcohol oxidase activity was calculated with the formula

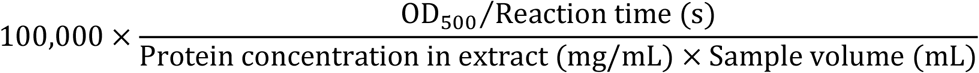

and reported in units mgp^−1^.

### Calculating substrate consumption and protein expression rates from the data

We are concerned with experiments in which a chemostat is fed with the primary carbon source *S*_1_ (methanol) and a secondary carbon source *S*_2_ which may be repressing (glycerol) or non-repressing (sorbitol). The primary carbon source *S*_1_ induces the synthesis of the enzyme *E*_1_ which represents LacZ or AOX since the latter is expressed almost entirely from an *AOX1* promoter. We are interested in measuring the steady state concentrations of biomass *X*, primary carbon source *S*_1_, and secondary carbon source *S*_2_ as well as the specific activity of enzyme *E*_1_. These quantities are denoted *x*, *s*_1_, *s*_2_, and *e*_1_, respectively, and satisfy the mass balances:

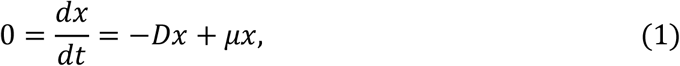

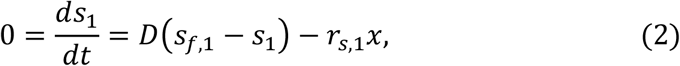

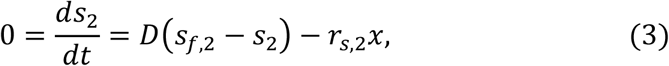

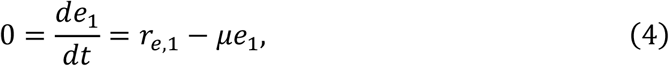

where *s*_*f*,1_, *s*_*f*,2_ denote the respective feed concentrations of *S*_1_, S_2_; and *μ*, *r*_*s*,1_, *r*_*s*,2_, *r*_*e*,1_ denote the respective specific rates of growth, consumption of substrate, and expression of a stable intracellular protein (Pfeffer *et* al., 2011; Singh and Narang, 2020). It follows from Eqs. (1)–(4) that

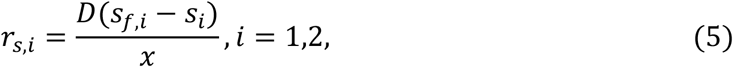

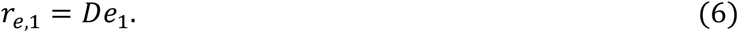

These equations were used to calculate *r*_*s*,1_, *r*_*s*,2_, and *r*_*e*,1_ from the measured values of the operating conditions *D*, *s*_*f*,*i*_ and the steady state concentrations *s*_*i*_, *x*, and *e*_1_.

## Results

### Substrate consumption and P_AOX1_ expression in the presence of glycerol and sorbitol

Our goal is to study the kinetics of substrate consumption and P_*AOX1*_ expression during mixed-substrate growth on methanol + glycerol and methanol + sorbitol; however, we also characterized the substrate consumption kinetics during single-substrate growth on glycerol and sorbitol. In batch (shake-flask) cultures grown on glycerol and sorbitol, the biomass yields were quite similar (~0.6 gdw g^−1^), but the maximum specific growth rates *μ*_m_ were dramatically different (Table 1). Due to the exceptionally small *μ*_*m*_ of 0.03 h^−1^ on sorbitol, we could not perform chemostat experiments with pure sorbitol, but we did perform such experiments with glycerol. We found that the biomass and residual glycerol concentrations followed the pattern characteristic of single-substrate growth in continuous cultures (Fig. 1a). The specific glycerol consumption rate, calculated from these data using Eq. (5), increased linearly with *D* with a significant positive *y*-intercept (Fig. 1b). Fitting these data to Pirt’s model (Pirt, 1965) gave a true biomass yield of 0.67 gdw g^−1^, and specific maintenance rate of 0.07 g gdw^−1^ h^−1^. The specific LacZ and AOX activities which were positively correlated in general, decreased with *D* (Fig. 1c). The specific LacZ and AOX expression rates, calculated from the data in Fig. 1c using Eq. (6), did not exceed ~1000 and ~300 units mgp^−1^ h^−1^, respectively (Fig. 1d).

**Table 1:** Maximum specific growth rates and biomass yields during single-substrate growth of the Mut^+^ strain of *K. phaffii* on glycerol and sorbitol. The true biomass yield in the chemostat was determined by fitting the variation of the specific substrate consumption rate with *D* to Pirt’s model.

| Carbon source | Maximum specific growth rate (h<sup>-1</sup>) | Biomass yield in shake flask (gdw g<sup>-1</sup>) | True biomass yield in chemostat (gdw g<sup>-1</sup>) |
| --- | --- | --- | --- |
| Glycerol | 0.24 ± 0.01 | 0.61 ± 0.03 | 0.67 |
| Sorbitol | 0.03 ± 0.01 | 0.56 ± 0.01 | ND |

**Fig. 1:**
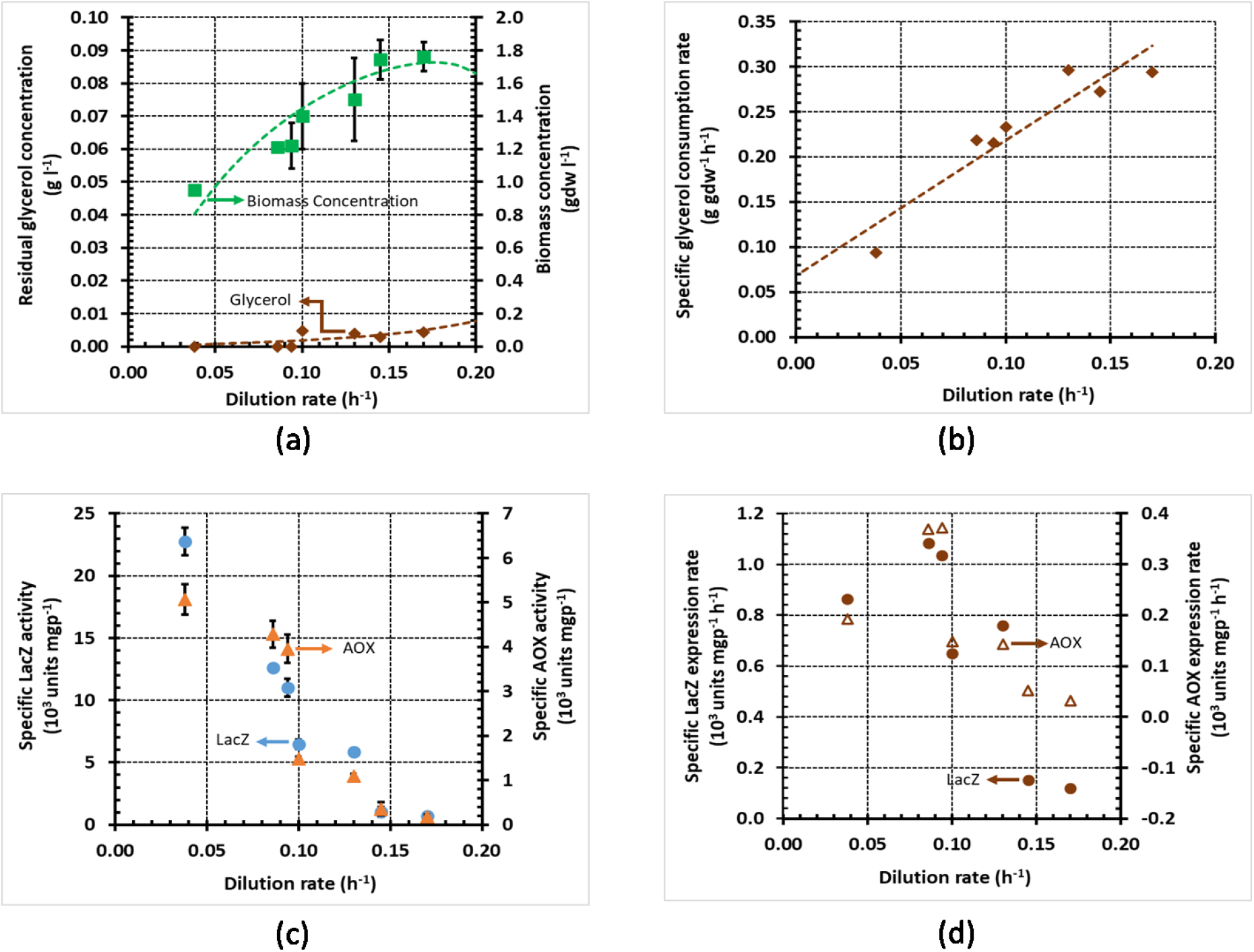
Variation of steady state concentrations and rates with the dilution rate during growth of *K. phaffii* strain Mut^+^ (pSAOH5-T1) in a chemostat fed with glycerol (~3.1 g l^−1^). (a) Concentrations of biomass and residual glycerol. (b) Specific glycerol consumption rates calculated from the data in (a) using Eq. (5). (c) Specific activities of LacZ and AOX. (d) Specific Lac Z and AOX expression rates calculated from the data in (c) using Eq. (6).

### Substrate consumption and P_AOX1_ expression in the presence of mixtures

When the Mut^+^ strain is grown in batch cultures of methanol + glycerol and methanol + sorbitol, there is diauxic growth, but methanol is the *unpreferred* substrate during growth on methanol + glycerol, and the *preferred* substrate during growth on methanol + sorbitol (Ramón *et al*., 2007). Such mixtures, which display diauxic growth in batch cultures, exhibit a characteristic substrate concentration profile in continuous cultures (Egli *et al*., 1986; Noel and Narang, 2009) (Supplementary Fig. S1a). In the *dual-limited* regime, which extends up to dilution rates approximately equal to the *μ*_*m*_ for the unpreferred substrate, both substrates limit growth because their residual concentrations *s*_*i*_ are on the order of their saturation constants *K*_*s*,*i*_ (*s*_*i*_~*K*_*s*,*i*_), and therefore, both substrates are completely consumed (*s*_*i*_ ≪ *s*_*f*,*i*_). Beyond the dual-limited regime, only the preferred substrate limits growth because the residual concentration of the unpreferred substrate is well above its saturation constant. At the intermediate *D* corresponding to the *transition* regime, the preferred substrate is still consumed completely, but the unpreferred substrate is only partially consumed. Beyond the transition regime, the unpreferred substrate is not consumed at all.

When methanol + glycerol and methanol + sorbitol were fed to a continuous culture, the variation of the substrate concentrations with *D* was consistent with the characteristic pattern described above. In the dual-limited regime, both substrates were completely consumed — up to *D* = 0.08 h^−1^ ≈ 0.11 h^−1^ = *μ*_*m*_|_methanol_ (Singh and Narang, 2020) in Fig. 2a and *D* = 0.03 h^−1^ = *μ*_*m*_|_sorbitol_ h^−1^ in Fig. 3a. In the transition regime, the unpreferred substrate was partially consumed up to dilution rates well above its *μ*_*m*_ — up to *D* = 0.2 ≈ 2 × *μ*_*m*_|_methanol_ h^−1^ in Fig. 2a, and up to *D* = 0.08 ≈ 3 × *μ*_*m*_|_sorbitol_ h^−1^ in Fig. 3a.

**Fig. 2:**
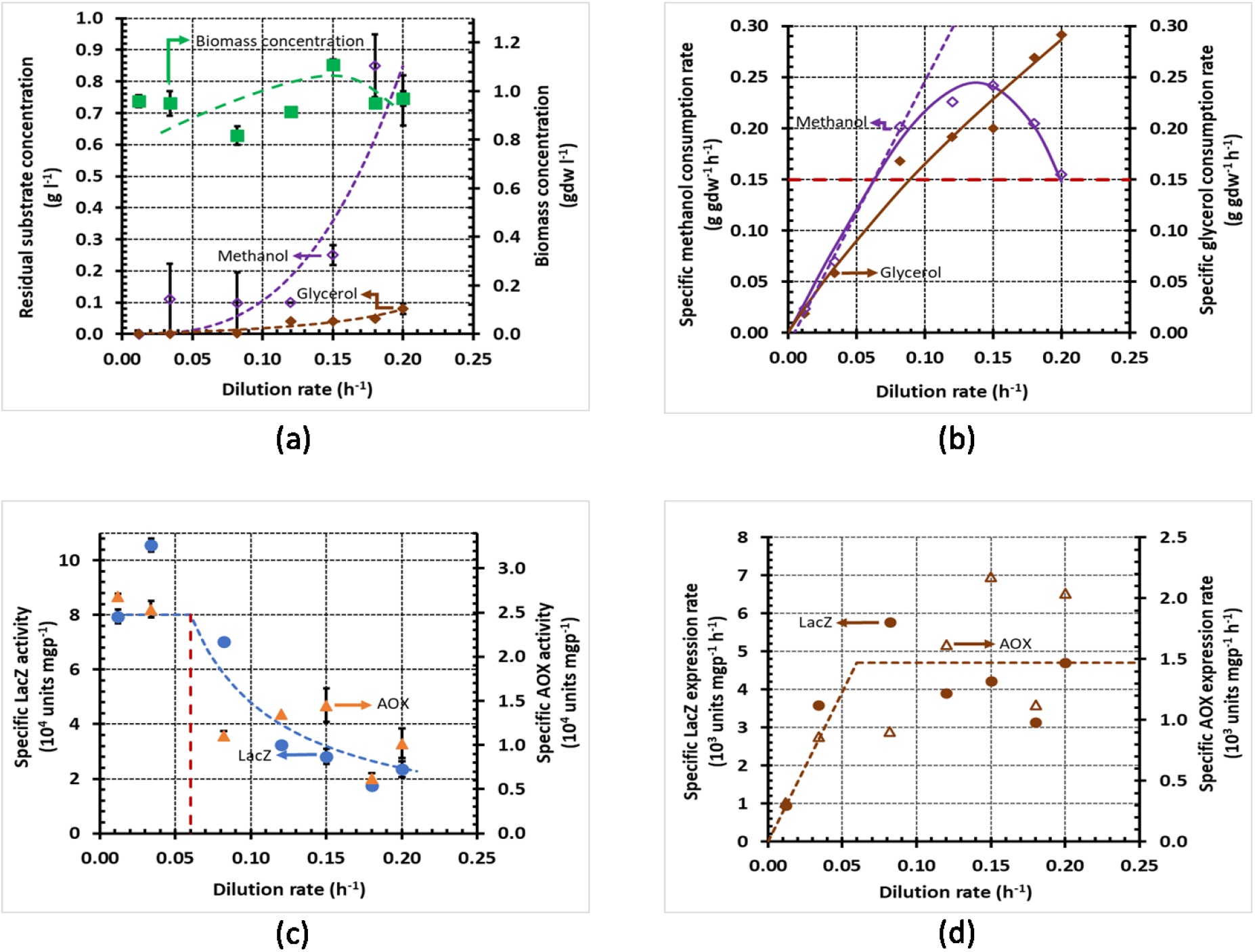
Variation of steady state concentrations with the dilution rate during growth of *K. phaffii* strain Mut^+^ (pSAOH5-T1) in a chemostat fed with a mixture of glycerol (~1.5 g l^−1^) and methanol (~1.6 g l^−1^). (a) Concentrations of biomass, residual glycerol, and residual methanol (b) Specific methanol and glycerol consumption rates calculated from the data in (a) using Eq. (5). The dashed line passing through the origin shows the linear increase of the specific methanol consumption rate in the dual-limited regime. The horizontal dashed line shows the threshold specific methanol consumption rate of 0.15 g gdw^−1^ h^−1^. (c) Specific activities of LacZ and AOX. (d) Specific LacZ and AOX expression rates calculated from the data in (c) using Eq. (6).

**Fig. 3:**
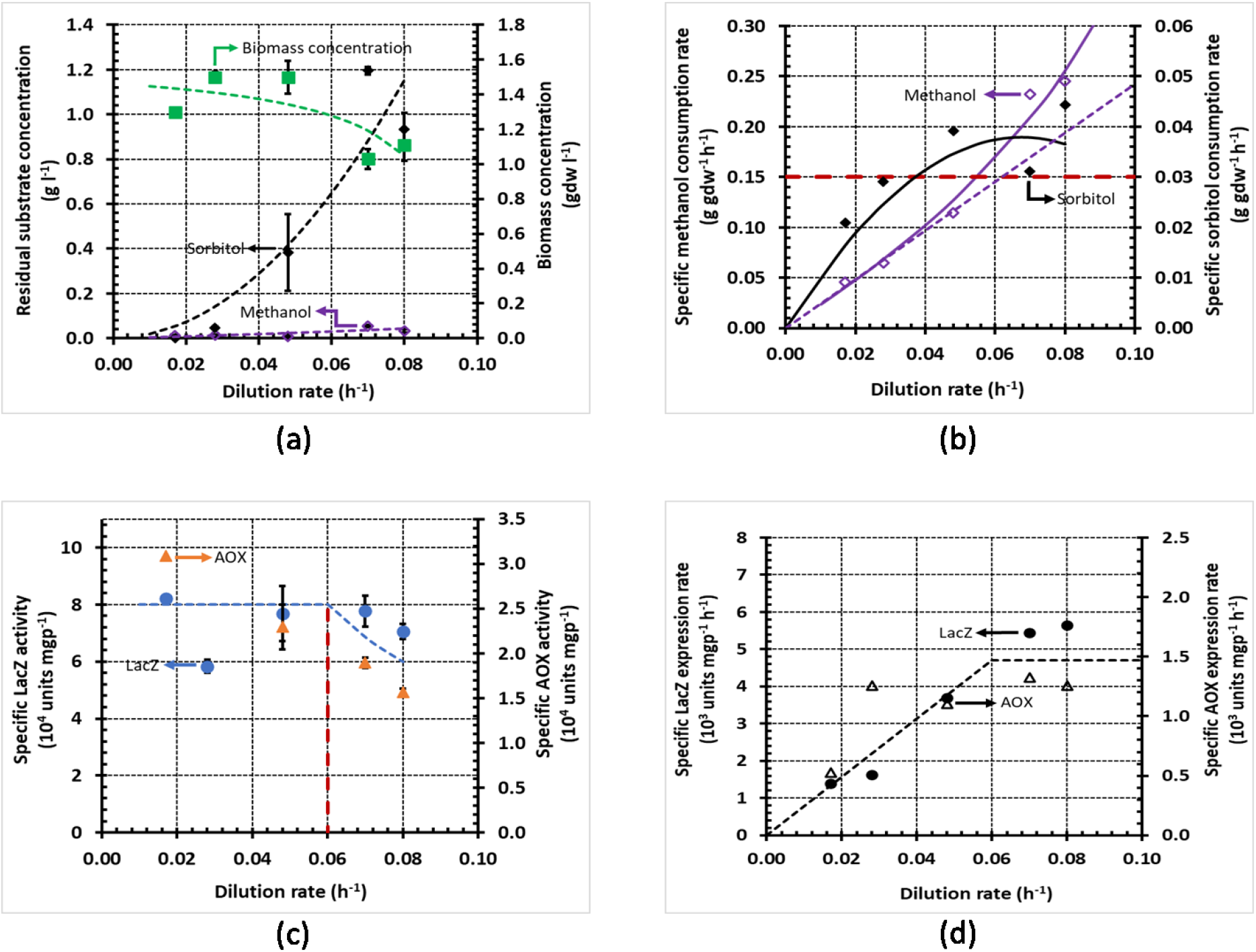
Variation of steady state concentrations with the dilution rate during growth of *K. phaffii* strain Mut^+^ (pSAOH5-T1) in a chemostat fed with a mixture of sorbitol (~1.5 g l^−1^) and methanol (~3.2 g l^−1^). (a) Concentrations of biomass, residual sorbitol and residual methanol. (b) Specific methanol and glycerol consumption rates calculated from the data in (a) using Eq. (5). The dashed line passing through the origin shows the linear increase of the specific methanol consumption rate in the dual-limited regime. The horizontal dashed line shows the threshold specific methanol consumption rate of 0.15 g gdw^−1^ h^−1^. (c) Specific activities of LacZ and AOX. (d) Specific LacZ and AOX expression rates calculated from the data in (c) using Eq. (6).

During single-substrate growth, the specific substrate consumption rate usually increases linearly with *D* up to washout (Pirt, 1965), but during mixed-substrate growth, the specific substrate consumption rates increase linearly with *D* only in the dual-limited regime (Egli *et al*., 1986; Noel and Narang, 2009) (Supplementary Fig. S1b). The dashed lines in Figs. 2b and 3b show that during growth on methanol + glycerol and methanol + sorbitol, the specific methanol consumption rate is indeed proportional to *D* up to *D* = 0.08 h^−1^ and *D* = 0.03 h^−1^, respectively. Beyond the respective dual-limited regimes, the specific methanol consumption rates change non-linearly (Supplementary Fig. S1b). In the case of methanol + glycerol, the specific methanol consumption rate decreases nonlinearly beyond *D* = 0.08 h^−1^ due to repression of methanol consumption by glycerol (Fig. 2b); in the case of methanol + sorbitol, the specific methanol consumption rate increases non-linearly beyond *D* = 0.03 h^−1^ due to the enhanced methanol consumption that occurs to compensate for repression of sorbitol consumption by methanol (Fig. 3b). Now, by a judicious choice of the feed concentrations calculated from Egli’s model for dual-limited growth (Egli et al., 1993), we ensured that when growth on both the mixtures is dual-limited (*D* ≤ 0.03 h^−1^), the specific methanol consumption rates of the two mixtures are not only proportional to *D*, but also equal in magnitude. The specific methanol consumption rates of the two mixtures start diverging beyond *D* = 0.03 h^−1^, but they remain approximately equal up to *D* = 0.05 h^−1^.

Although it is widely accepted that glycerol is repressing and sorbitol is non-repressing in batch cultures, we found remarkably similar specific LacZ and AOX activities and expression rates in continuous cultures fed with methanol + glycerol and methanol + sorbitol. At low dilution rates (*D* ≤ 0.05 h^−1^), when both mixtures support equal specific methanol consumption rates, the specific LacZ and AOX activities on both mixtures are also equal (Figs. 2c and 3c), and hence, their specific LacZ and AOX expression rates are also the same (Figs. 2d and 3d). At high dilution rates (*D* ≥ 0.05 h^−1^), the specific methanol consumption rates of both mixtures change substantially, but the specific LacZ and AOX expression rates are relatively insensitive to this change. Indeed, in the case of methanol + glycerol, the specific methanol consumption rate doubles when *D* increases from 0.05 h^−1^ to 0.12 h^−1^, and decreases 40 % when *D* increases from 0.12 h^−1^ to 0.20 h^−1^. But the specific LacZ and AOX activities decrease inversely with *D* (Fig. 2c), and hence, the specific LacZ and AOX expression rates are constant (Figs. 2c and 2d). In the case of methanol + sorbitol, the specific methanol consumption rate doubles when *D* increases from 0.05 h^−1^ to 0.08 h^−1^, but the specific LacZ and AOX expression rates increase only 25 % (Fig. 3d). Furthermore, the constant maximum specific LacZ and AOX expression rates of 4000–6000 units mgp^−1^ h^−1^ and 1200–2000 units mgp^−1^ h^−1^, respectively, are close to the corresponding maximum values observed during growth on methanol + glycerol. Taken together, these data suggest that the specific P_*AOX1*_ expression rate is a function (i.e., completely determined by) the specific methanol consumption rate.

The specific P_AOX1_ expression rate is a function of the specific methanol consumption rate To test this hypothesis, we plotted the specific LacZ and AOX expression rates *r*_*e*,1_ at various *D* in Figs. 2d–3d against the corresponding specific methanol consumption rate *r*_*s*,1_ in Figs. 2b–3b. This yielded the graph in Fig. 4 which shows that at every specific methanol consumption rate, both mixed-substrate cultures have approximately the same specific P_*AOX1*_ expression rate. The specific P_*AOX1*_ expression rate is therefore completely determined by the specific methanol consumption rate regardless of the type (repressing or non-repressing) of the secondary carbon source. More precisely, the specific P_*AOX1*_ expression rate, *r*_*e*,1_ is proportional to the specific methanol consumption rate, *r*_*s*,1_ up to the threshold value ~0.15 g gdw^−1^ h^−1^ and remains approximately constant thereafter at the maximum value of ~5 units gdw^−1^ h^−1^. Hence, the specific P_*AOX1*_ expression rates of the mixtures can be approximated by the piecewise linear function

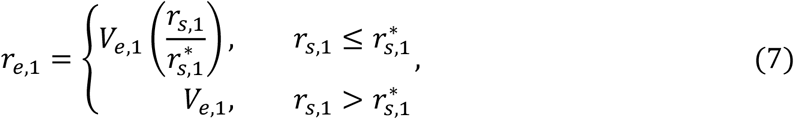

**Fig. 4:**
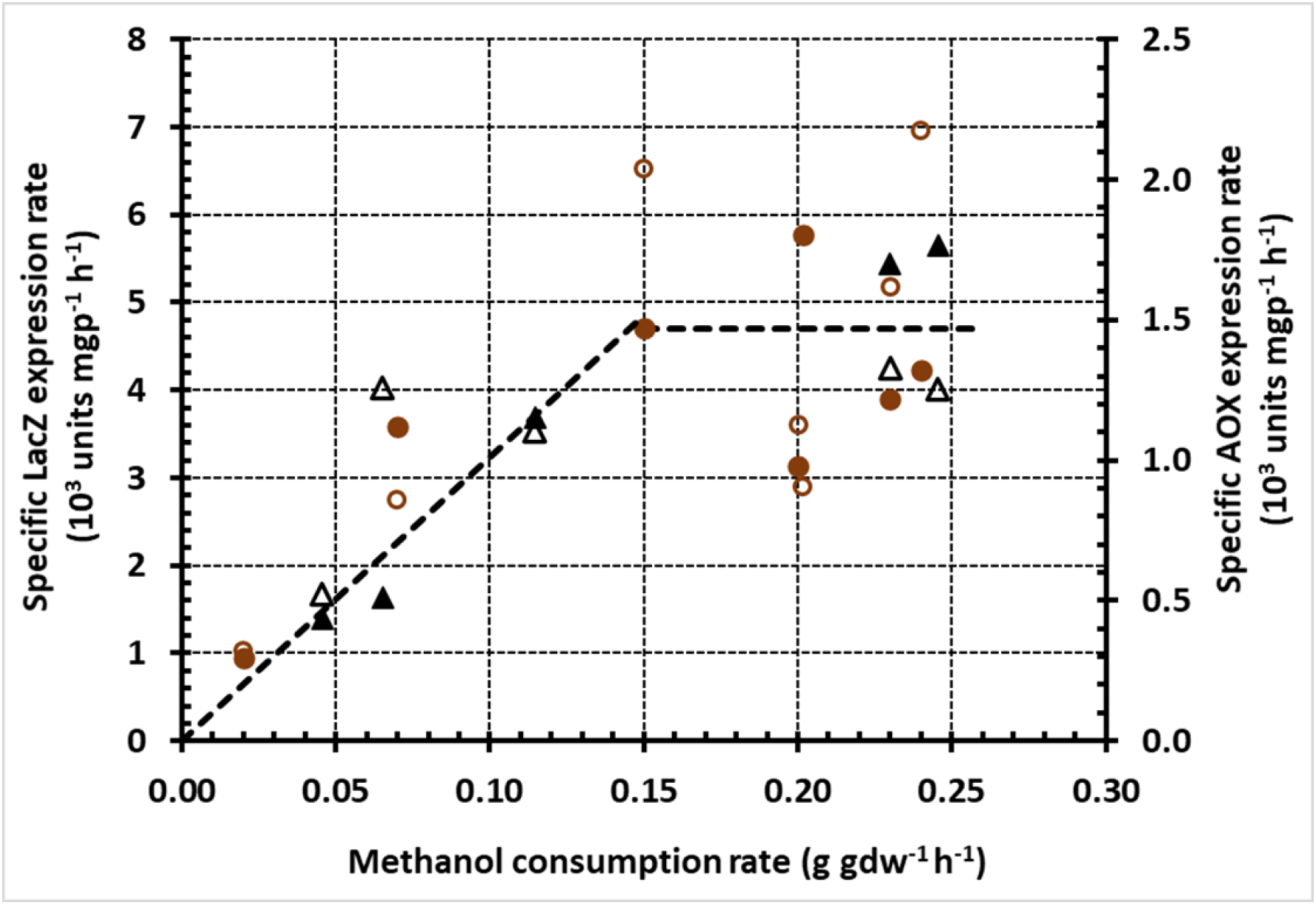
Variation of the specific LacZ (closed symbols) and AOX (open symbols) expression rates with the specific methanol consumption rate during growth on methanol + glycerol (brown circles) and methanol + sorbitol (black triangles) The graph was obtained by plotting the specific methanol consumption rates in Figs. 2b–3b against the corresponding specific LacZ and AOX expression rates in Figs. 2d–3d.

where *V*_*e*,1_ denotes the maximum specific P_*AOX1*_ expression rate, and *r*^*^_*s,1*_ denotes the threshold specific methanol consumption rate beyond which the specific P_*AOX1*_ expression rate has its maximum value *V*_*e*,1_.

## Discussion

Our main conclusion is that over the range of dilution rates considered in our work (0.02–0.2 h^−1^), the P_*AOX1*_ expression rate is completely determined by the methanol consumption rate regardless of the type of the secondary carbon source. This conclusion may appear to subvert the prevailing consensus according to which the expression rate of a promoter is strongly inhibited in the presence of repressing secondary carbon sources. However, this conclusion is based on studies with *batch* cultures. We show below that our conclusion is consistent with the *continuous* culture studies reporting the expression of not only the *AOX1* promoter of *K. phaffii* but also the exemplary *lac* promoter of *E. coli*.

### Comparison with chemostat studies of P_*AOX1*_ expression by *K. phaffii*

Jungo *et al* performed their mixed-substrate studies by fixing *D*, *s*_*f*,1_ + *s*_*f*,2_ and increasing the fraction of methanol in the feed *σ*_1_ = *s*_*f*,1_⁄(*s*_*f*,1_ + *s*_*f*,2_) at a slow linear rate aimed at maintaining quasi-steady state. They found that as *σ*_1_ increased:

a. The residual methanol remained negligibly small, and the biomass concentration decreased linearly.
b. The specific biotin expression rate increased hyperbolically until it reached a maximum, which was essentially the same for both mixtures.

It follows from a) that the specific methanol consumption rate, which is approximately equal to *D*(*s*_*f*,1_ + *s*_*f*,2_)*σ*_1_⁄*x*, increased throughout their experiment. But then b) implies that as the specific methanol consumption rate increased, the specific biotin expression rate of all three cultures reached essentially the *same* maximum (cf. Fig. 5).

Berrios and co-workers compared the methanol consumption and ROL production rates of the Mut^+^ strain at two different temperatures (22 and 30 °C) during growth on methanol, methanol + glycerol, and methanol + sorbitol (Berrios *et al*., 2017). These experiments were done in chemostats operated at *D* = 0.03 h^−1^, and in the case of mixed-substrate experiments, fed with two feed compositions (40 and 70 C-mole % methanol). They found that, “Sorbitol-based cultures led to a higher *q*_*p*_ than both glycerol-based and control cultures at most studied conditions.” But, closer inspection shows that that in all their experiments, the specific expression rates were 0.8–0.9 units gdw^−1^ h^−1^, which is close to the maximum specific expression rate of 1 unit gdw^−1^ h^−1^. It is therefore conceivable that the higher productivities observed with sorbitol-based cultures are not statistically significant.

### Comparison with chemostat studies of expression by *lac* promoter of *E. coli*

Analogous results have also been obtained in studies of *lac* expression in *E. coli*. Indeed, batch experiments with mixtures of lactose + glycerol, lactose + glucose, and lactose + glucose-6-phophate show that glycerol is non-repressing, whereas glucose and glucose-6-phosphate are repressing (Magasanik, 1970). However, when chemostat experiments were performed with these three mixtures (Smith and Atkinson, 1980), they yielded the *same* steady state specific β-galactosidase (LacZ) activity at all *D* ≲ 0.5 h^−1^ (Supplementary Fig. S2). Furthermore, when the steady state specific LacZ activities at various *D* were plotted against the corresponding specific lactose consumption rates at the same *D*, the data for all three mixtures collapsed into a single line (Supplementary Fig. S3). This led the authors to conclude that the steady state specific LacZ activity was “an apparently linear function of the rate of lactose utilization independent of the rate of metabolism of substrates other than lactose which are being concurrently utilized.” But then it follows from Eq. (6) that the steady state specific LacZ expression rate is also completely determined by the specific lactose consumption rate regardless of the type (repressing or non-repressing) of the secondary carbon source (Supplementary Fig. S4).

In conclusion, the specific P_*AOX1*_ expression rate of *K. phaffii* appears to be completely determined by the specific methanol consumption rate regardless of the type (repressing or non-repressing) of the secondary carbon source. Analysis of the literature shows that the specific expression rate of the *lac* operon of *E. coli* is also completely determined by the specific lactose consumption rate regardless of the type of secondary carbon source. It would be interesting to explore if similar results are obtained in other microorganisms and substrate mixtures.

## Declarations

### Funding Information

The authors would like to thank Department of Biotechnology (Government of India) for funding this project (grant BT/PR13831/BBE/117/68/2015).

### Conflict of Interest

The authors declare that they have no conflict of interest.

### Availability of data and material

Raw data is available upon request.

### Code availability

Not applicable

### Author’s contribution

AS and AN conceived and designed the research. AS conducted the experiments. AS and AN analysed the data and wrote the manuscript. All authors read and approved the manuscript.

### Compliance with ethical standards

The authors declare that no human participants or animals were used for the purpose of this study.

### Consent to participate

Not applicable

## Notes

Funding Information: This research was supported by the grant BT/PR13831/BBE/117/68/2015 received from the Department of Biotechnology (DBT), Government of India.

### Competing Interest Statement

The authors have declared no competing interest.

### Summary of Updates

The manuscript has been updated to replace mathematical equations with verbal explanations.

