## Supplementary Data for "P_*AOX1*_ expression in mixed-substrate continuous cultures of *Komagataella phaffii* (*Pichia pastoris*) is completely determined by methanol consumption regardless of the secondary carbon source"

**Running title: P*_AOX1_* expression is completely determined by methanol consumption**

Anamika Singh^1,2^ and Atul Narang^1,^*

^1^Department of Biochemical Engineering and Biotechnology, Indian Institute of Technology, Hauz Khas, New Delhi 110016, India

^2^Present address: International Centre for Genetic Engineering and Biotechnology, New Delhi 110067, India

Funding Information: This research was supported by the grant BT/PR13831/BBE/117/68/2015 received from the Department of Biotechnology (DBT), Government of India.

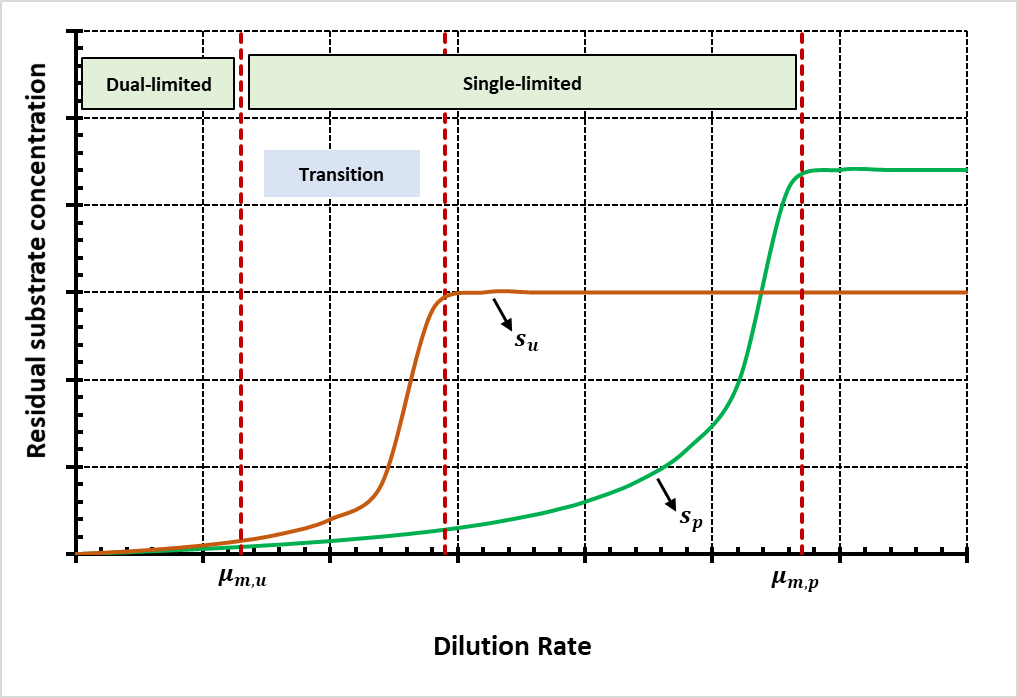

(a)

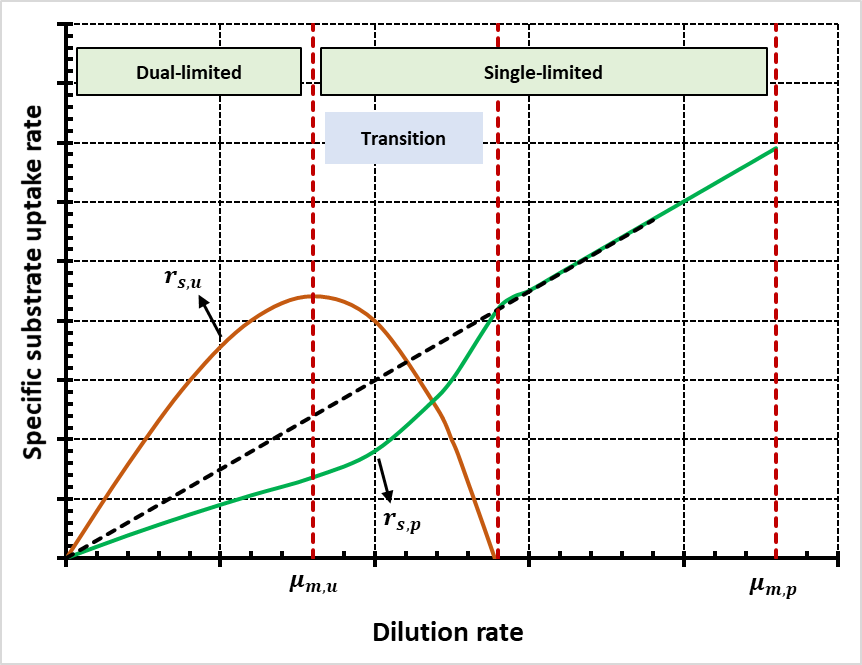

(b)

**Fig. S1:** Schematic representation of the substrate concentration and uptake rate patterns observed in continuous cultures fed with mixtures that exhibit diauxic growth in batch cultures (adapted from Egli *et al.*, 1986 and Noel and Narang, 2009). The symbols $\mu_{m,p}$ and $\mu_{m,u}$ denote the maximum specific growth rates on the preferred and unpreferred substrates during diauxic growth. (a) The variation with $D$ of the residual concentration of the preferred substrate $s_{p}$ and the unpreferred substrate $s_{u}$. (b) The variation with $D$ of the specific uptake rate of the preferred substrate $r_{s,p}$ and the unpreferred substrate $r_{s,u}$.

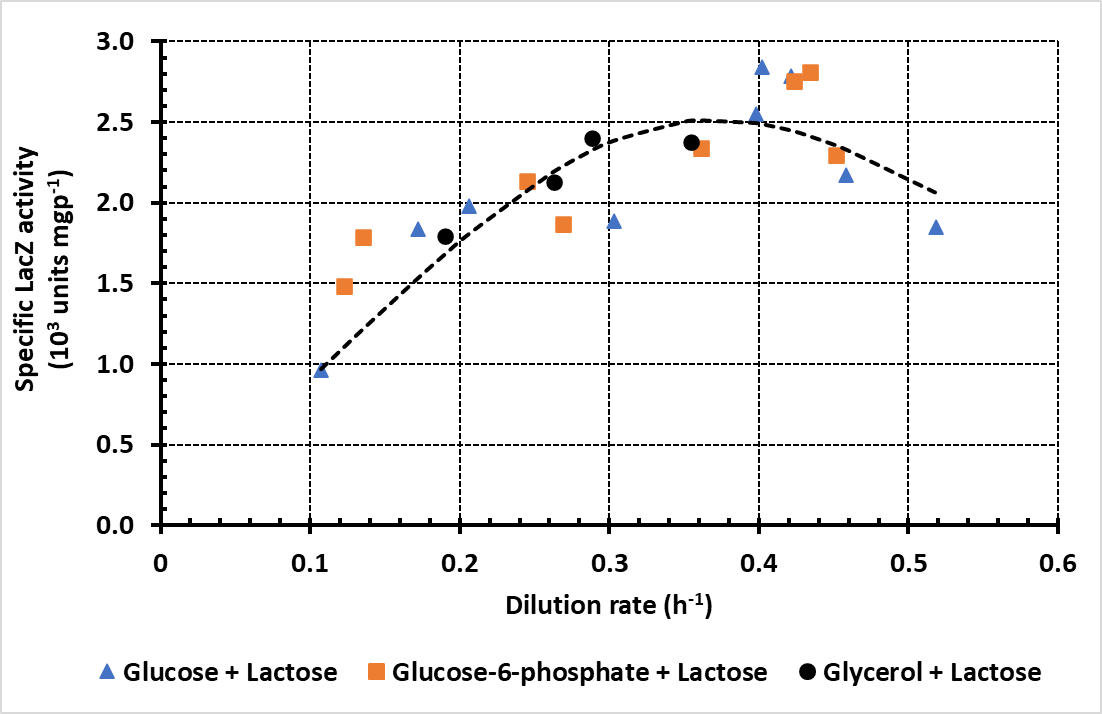

**Fig. S2:** Variation of the steady state specific LacZ activity with the dilution rate during growth of *E. coli* in chemostats fed with mixtures of lactose (1 mM) + glucose (2mM), lactose (1 mM) + glucose-6-phosphate (2mM), and lactose (1 mM) + glycerol (4 mM), as reported in Fig. 3 of Smith and Atkinson, 1980.

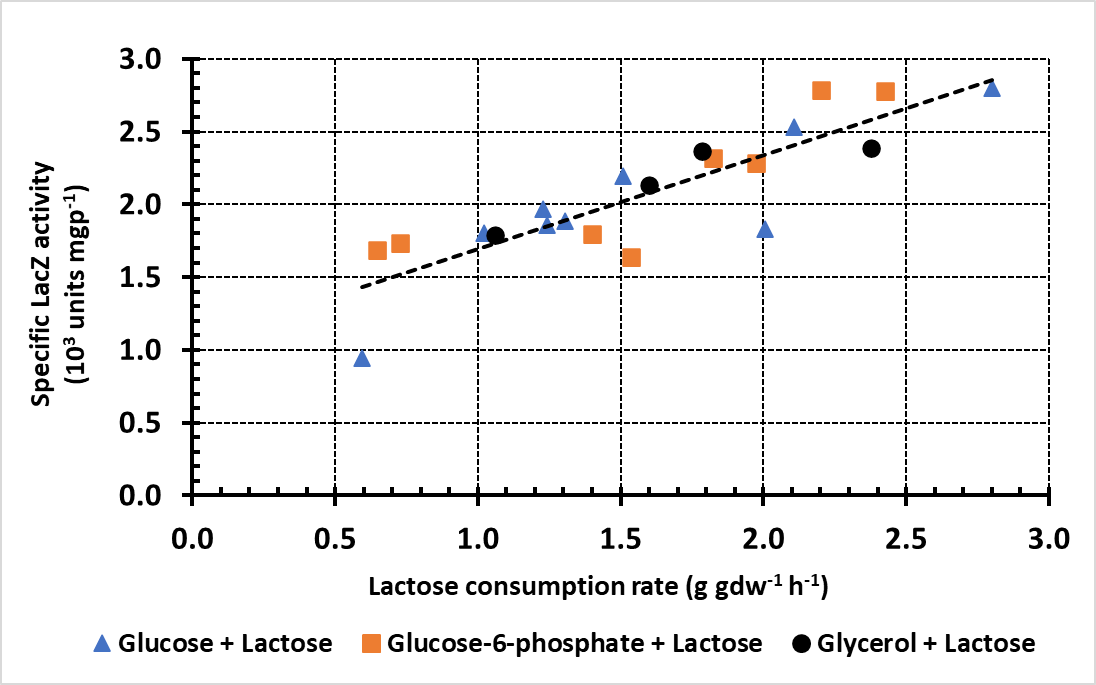

**Fig. S3:** Variation of the steady state specific LacZ activity at various dilution rates with the specific lactose consumption rate at that dilution rate during growth of *E. coli* in chemostats fed with mixtures of lactose (1 mM) + glucose (2mM), lactose (1 mM) + glucose-6-phosphate (2mM), and lactose (1 mM) + glycerol (4 mM), as reported in Fig. 6 of Smith and Atkinson, 1980.

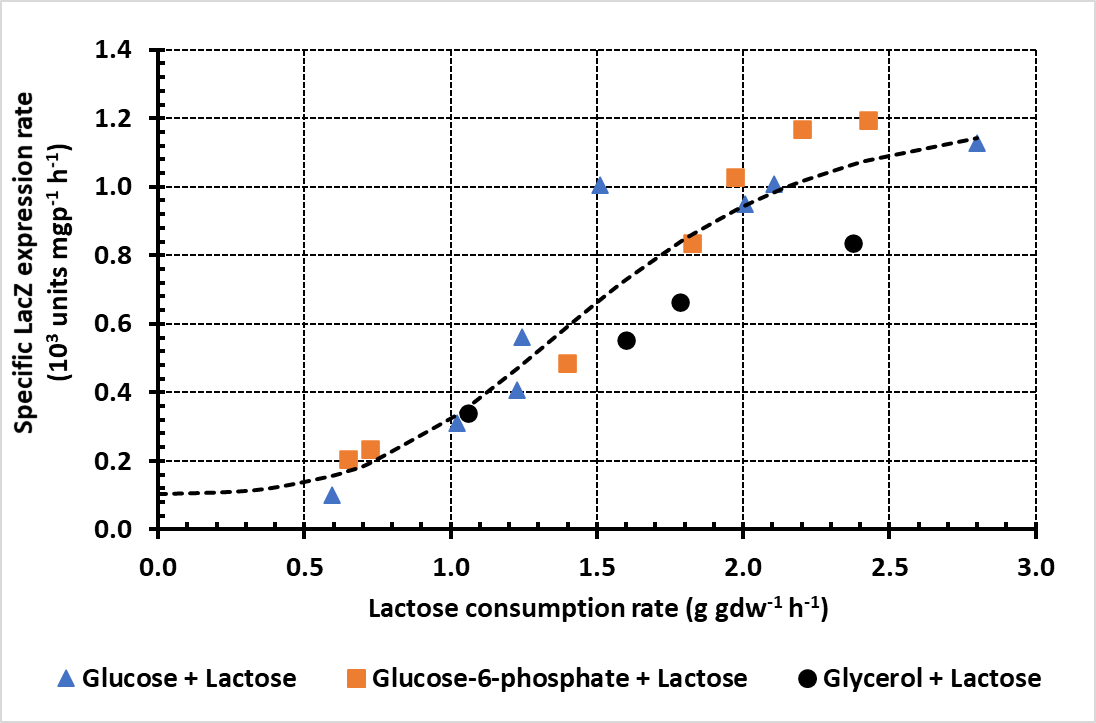

**Fig. S4:** Variation of the steady state specific LacZ expression rate at various dilution rates with the specific lactose consumption rate at that dilution rate during growth of *E. coli* in chemostats fed with mixtures of lactose (1 mM) + glucose (2mM), lactose (1 mM) + glucose-6-phosphate (2mM), and lactose (1 mM) + glycerol (4 mM). The data were derived from Fig. S3 by using Eq. (6) to convert the specific LacZ activities to specific LacZ expression rates.
